# Tau pathology in early Alzheimer’s disease disrupts selective neurophysiological network dynamics

**DOI:** 10.1101/524355

**Authors:** Ece Kocagoncu, Andrew Quinn, Azadeh Firouzian, Elisa Cooper, Andrea Greve, Roger Gunn, Gary Green, Mark W. Woolrich, Richard N. Henson, Simon Lovestone, Deep and Frequent Phenotyping study team, James B. Rowe

**Author notes:** Correspondence to: Ece Kocagoncu, Cambridge Centre for Frontotemporal Dementia and Related Disorders, University of Cambridge, Department of Clinical Neurosciences, Herchel Smith Building, Forvie Site, Robinson Way, Cambridge Biomedical Campus, Cambridge, CB2 0SZ, UK.

## Abstract

The role of aggregation of misfolded Tau protein in the pathogenesis of Alzheimer’s disease is the subject of rapid biomarker development and new therapeutic strategies to slow or prevent dementia. We tested the hypothesis that Tau pathology is associated with functional organization of widespread neurophysiological networks. We used electro-magnetoencephalography (E/MEG) in combination with [^18^F]AV1451 PET scanning to quantify Tau-dependent network disruption. Using a graph theoretical approach to MEG connectivity, we quantified nodal measures of functional segregation, centrality and efficiency of information transfer. We correlated these metrics against the nodes’ uptake of [^18^F]AV1451. There were both regional- and frequency-specific effects of Tau levels on the efficiency of information transfer and network segregation in early AD. Tau correlated with temporal regional participation coefficient (in delta, theta, beta bands); and temporal lobar eigenvector centrality (in theta, alpha, beta bands), but greater eccentricity at higher frequencies (gamma). The results support the translational development of neurophysiological “signatures” as biomarkers of Alzheimer’s disease, with potential to facilitate experimental medicines studies.

## 1. Introduction

There is a pressing need for new therapeutic strategies to prevent or arrest Alzheimer’s disease, especially where applicable at the prodromal stage of disease. The evaluation of new candidate compounds requires robust tools that are sensitive to the pathogenic mechanisms in early stages of disease. Commonly used tools to quantify the effects of Alzheimer’s disease pathology in people vary in the degree of invasiveness (e.g. magnetic resonance imaging *versus* lumbar puncture), cost and scalability for large trials (e.g. blood tests *versus* positron emission tomography), and the degree to which they provide mechanistic insight into the pathogenesis of Alzheimer’s disease (e.g. cognitive tests *versus* tau-ligand PET).

In this study, we assess a neurophysiological perspective that links recent advances in preclinical and translational models of Alzheimer’s disease. There are two key aspects to our approach. First, is the recognition of the effect of Tau and Aβ on synaptic dysfunction (Ittner et al., 2010; Murray et al., 2015), early in the cascade of Alzheimer pathogenesis and without atrophy or cell death. This in turn impairs the network dynamics, which underpin cognition (Ahmed et al., 2014; Kimura et al., 2014). Tau and Aβ induced changes in GABAergic function (Li, G. et al., 2009), and glutamatergic function (Hsieh et al., 2006; LaFerla and Oddo, 2005; Li, S. et al., 2009; Liu et al., 2004; Shankar et al., 2007) will further disrupt effective communication in local and large scale neurocognitive networks.

Second, is the recognition of neurophysiological signatures of Alzheimer’s disease. For example, magnetoencephalograhy (MEG) distinguishes Alzheimer’s disease pathology from frontotemporal lobar degeneration by their spectral signatures, while retaining functional anatomical concordance with the clinical syndromes (Sami et al., 2018). The brain’s evoked and induced responses as recorded by MEG and electroencephalograhy (EEG) distinguish Alzheimer’s disease from controls, in advanced disease (Dauwels et al., 2010; Sitnikova et al., 2018), mild cognitive impairment stage, and even pre-symptomatically in carriers of autosomal dominant mutations (Ochoa et al., 2017; Suarez-Revelo et al., 2016). The spectral features of non-invasive clinical studies recapitulate invasive and ex-vivo recordings of transgenic model systems (Koss et al., 2016; Kurudenkandy et al., 2014; Sami et al., 2018). Thus, MEG and EEG offer one way to capture synaptic dysfunction before extensive brain atrophy. However, the relationship between these physiological indices and the characteristic Tau pathology of human Alzheimer’s disease is unknown.

In the current study, we exploit the spatiotemporal precision of MEG to study network connectivity and oscillatory patterns, across different frequency bands. We use a graph theoretical approach, to extract regional and frequency specific summary measures of complex network function (Bullmore and Sporns, 2009). These graph metrics reflect the efficiency of information transfer and the extent to which a given region contributes to information processing at the modular or global level. For example, Alzheimer’s disease decreases the clustering coefficient and path length (i.e. measures of network efficiency at the local and global level) of network interactions in the alpha and beta bands (de Haan et al., 2009; Stam et al., 2009), while the reduction of small worldness (Vecchio et al., 2016) and eigenvector centrality of temporal areas (de Haan et al., 2012b) suggest an imbalance between local functional specialization and global integration.

We tested the relationship between such neurophysiological network changes and the progression of Tau pathology. Previous studies using fMRI based network connectivity measures (rather than MEG/EEG) have shown that the degree of connectivity of each cortical region correlates with expression of the MAPT gene for Tau (Rittman et al., 2016) and the accumulation of Tau as measured by [^18^F]AV1451 PET (Cope et al., 2018). This ligand binds to Tau aggregates in Alzheimer’s disease, in proportion to disease severity (Brier et al., 2016; Passamonti et al., 2017), and mirrors the distribution of pathology and functional deficits in variant presentations of Alzheimer’s disease (Ossenkoppele et al., 2016). We therefore used [^18^F]AV1451 PET to test the relationship between Tau pathology burden and MEG connectivity.

Our primary goal was to quantify the correlation of Tau burden with physiological network properties in early Alzheimer’s disease. A secondary goal was to measure the effect of Tau burden on the rate of change in these network properties, over six months. We hypothesized that in Alzheimer’s disease, (i) efficiency of information transfer at the local and global level is disrupted; (ii) the influence of central nodes on the network weaken; (iii) segregation of functional modules is reduced; (iv) and that these changes in graph metrics correlate with local increases in Tau burden across the cortex.

## 2. Materials and Methods

### 2.1 Study design

The Deep and Frequent Phenotyping study is a collaboration between the Dementias Platform UK and the NIHR Translational Research Collaboration in Dementia. It aims to assess the acceptability and feasibility of extensive and frequent phenotyping, to aid the design of larger scale future biomarker studies (Koychev et al., 2017). Here we report data from the pilot study phase, which included patients with early symptomatic Alzheimer’s disease. The study was approved by the National Research Ethics Committee London (REC reference 14/LO/1467). All participants had mental capacity and provided written informed consent.

### 2.2. Participants

12 participants with probable Alzheimer’s disease (McKhann et al., 1984) were recruited from local memory clinics (mean age: 69.94, age range: 54-82.7, 9 males, 3 females). Participants had a mean MMSE score of 24/30 (*SD =* 2.27), mean CDR score of 0.6/3 (*SD =* 0.31), and mean ADAS-Cog score of 13.9/70 (*SD =* 5.43) Participants had stable medication dose for at least a month, and for cholinesterase inhibitors and/or Memantine stable medication dose for at least three months.

### 2.3. MEG acquisition

MEG scans were acquired at rest (eyes open) over five minutes at four sites at baseline (Functional Imaging Laboratory at University College London, Oxford Centre for Human Brain Activity in University of Oxford, York Neuroimaging Centre at University of York, and MRC Cognition and Brain Sciences Unit at the University of Cambridge) using three types MEG scanners (CTF/VSM Omega 275, Elekta Vector View 306 and 4D Magnes 3600). Nine participants had a repeat E/MEG scan 6 months later.

Participants were seated in a magnetically shielded room and positioned under the MEG scanner in the upright position. EOG and ECG electrodes were used where available, plus head position indicator coils. For coregistration of the participant’s T1-weighted MRI scan to the MEG sensors, three fiducial points (nasion, left and right pre-auricular) and head surface points were digitized using Polhemus digitization. Simultaneous E/MEG was recorded continuously at 1000 Hz.

### 2.4. PET and MR

All PET scans were acquired at Imanova. MR scans were acquired at the Cambridge, Oxford and London sites, using Siemens 3T Trio with a 32-channel phased array head coil. 1 mm isotropic whole-brain structural 3D T1-weighted MPRAGE images were acquired using TI = 880 ms, TR = 2000 ms, and FA = 8° with a parallel imaging factor of 2. Two dynamic PET scans for Aβ and Tau were acquired on separate days. Participants were injected an intravenous bolus of [^18^F]AV1451 (120 min, 163 ± 10 MBq) and [^18^F]AV45 tracers (60 min, 150 ± 24 MBq) for Tau and Aβ respectively. A low dose CT scan immediately before each PET scan was used to estimate attenuation. The scans were acquired on Siemens PET/CT scanners (either Hi-Rez Biograph 6 or Biograph 6 TruePoint with TrueV, Siemens Healthcare, Erlangen, Germany). Dynamic images were reconstructed using a 2D Filtered Back Projection (FBP) algorithm resulting in a 128 × 128 matrix with 2 mm isotropic voxels. Corrections were applied for attenuation, randoms, scatter, and tracer radioactive decay.

Summary steps of the PET and MR preprocessing are given in Fig 1A (Firouzian et al., 2018). PET and MRI imaging processing was performed using MIAKAT™ (www.miakat.org). Each participant’s whole brain was extracted using the FMRIB software library (FSL) (Jenkinson et al., 2012), brain extraction tool (Smith, 2002) and the corresponding grey matter probability maps were created using SPM5 (www.fil.ion.ucl.ac.uk/spm). Further, dynamic PET data were corrected for motion. Regional time activity curves (TACs) were generated using the atlas and dynamic PET images. The simplified reference tissue model (SRTM) with cerebellar grey matter as a reference region were applied to the regional TACs to estimate the non-displaceable binding potential (BP_ND_) used to quantify the amount of tracer binding to the target proteins. The resulting BP_ND_ maps were coregistered to participant’s T1-weighted MRI scan. To correct for the partial volume effects, the Müller-Gärtner method was applied voxel-wise, which employs a 3-compartment model of the brain (i.e. white matter, grey matter and CSF tissue maps), as implemented in the PETPVE12 toolbox (Gonzalez-Escamilla et al., 2017). MR images and corrected PET images were normalized and resliced to match the atlas dimensions and resolution (1 mm isotropic). Current analysis focused on cortical Tau burden only.

**Fig 1.**
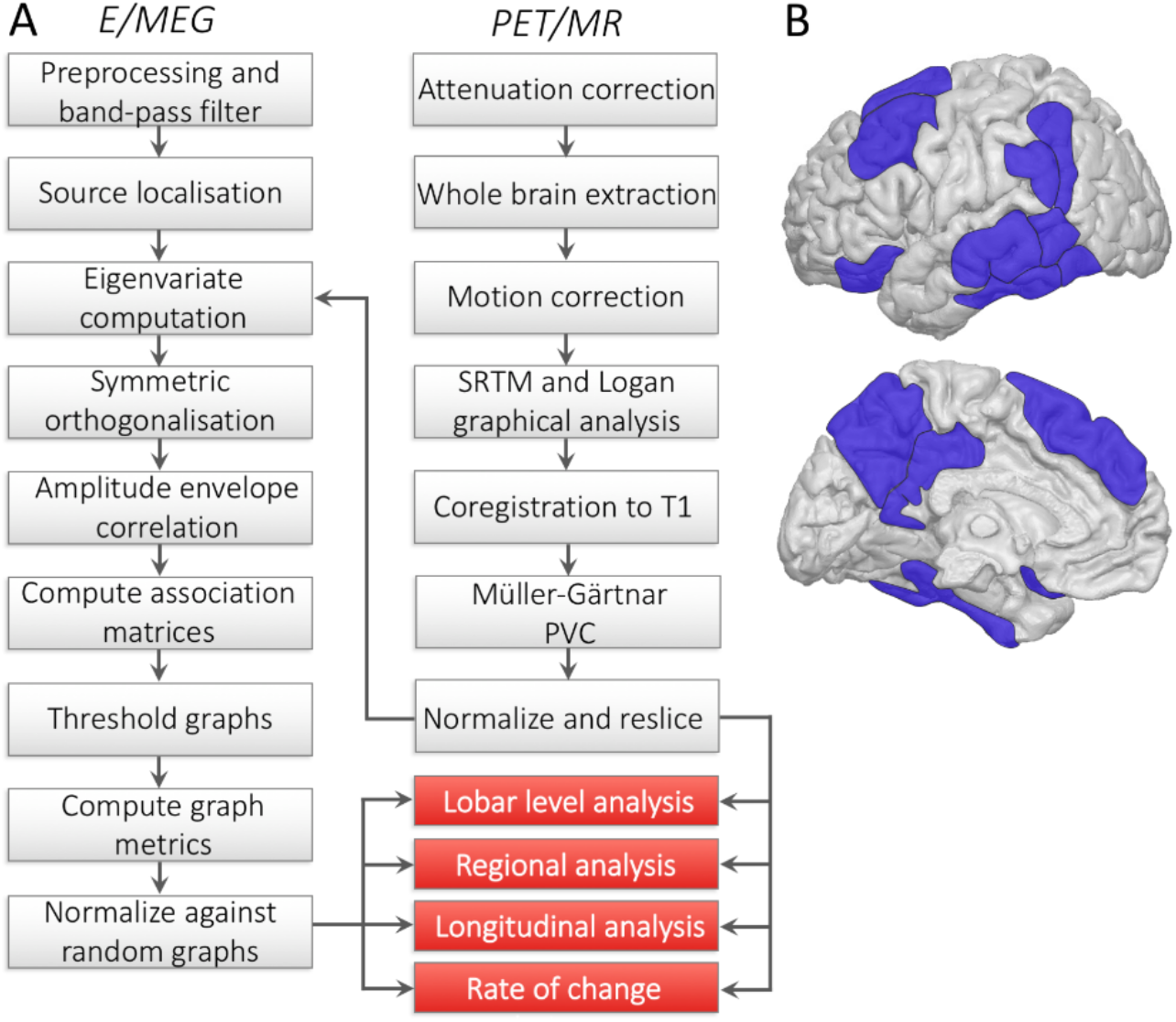
Details of the analysis. A. Pipeline showing the steps of E/MEG and PET/MR processing leading up to the graph theoretical analysis. B. Alzheimer’s disease related ROIs used in the regional analysis, subcortical areas not shown. Key: SRTM, simplified reference tissue model; PVC, partial volume correction.

### 2.5. E/MEG preprocessing and source localization

The raw MEG data acquired through Elekta scanners were pre-processed using MaxFilter 2.2 (Elekta Oy). Maxfiltering included detection and interpolation of bad sensors, signal space separation to remove external noise from the data and head movement correction. MEG data acquired through the CTF system were analyzed as third order synthetic gradiometers. Cardiac and blink artefacts were removed using an independent component analysis with 800 maximum steps and 64 principal components via the EEGLAB toolbox (Delorme and Makeig, 2004). On average 1.36 blink components (*SD =* 0.95) and 0.8 cardiac components (*SD =* 0.41) were removed. Summary steps of the E/MEG preprocessing are given in Fig 1A.

Data were further processed in SPM12 (www.fil.ion.ucl.ac.uk/spm). Data were bandpass filtered to five frequency bands of interest using fifth-order Butterworth filters: delta (0.1-4 Hz), theta (4-8 Hz), alpha (8-12 Hz), beta (12-30 Hz), and gamma (30-100 Hz). Data in the gamma band were further notch filtered to remove line noise. The continuous data were epoched into 4s long consecutive segments, resulting in approximately 75 epochs per participant. These segments were visually inspected for any remaining artefacts (e.g. motor) and bad channels and trials were removed. On average 5.46 (*SD =* 7.39) trials and 2.25 channels (*SD =* 2.75) were removed per participant. Data were then downsampled to 200 Hz.

The E/MEG data were source localized using all sensor types (Henson et al., 2009). The source space was modelled with a medium sized cortical mesh consisting of 8196 vertices via inverse normalization of SPM’s canonical meshes. Sensor positions were coregistered to the native T1-weighted MPRAGE scans using the fiducial and head shape points. Single shell and BEM models were used for forward modelling of MEG and EEG data respectively. Total power (induced and evoked)was estimated over the trials using the minimum norm estimate solution (R^2^ model fit: *M =* 91.49; *SD =* 6.56). Across all participants, the two visits and the frequency bands, five inversions showed R^2^ lower than 80%, and were excluded from the following analyses.

### 2.7. Graph theoretical analysis

A cortical graph was based on the Harvard Oxford atlas (HO) thresholded at 25%, with 98 cortical parcels including the hippocampi. The data were extracted from all the vertices that constitute each parcel, and their first eigenvariate was computed. Multivariate leakage correction method was applied that removes the zero lag effects across all parcels using symmetric orthogonalization (Colclough et al., 2015), allowing a more accurate estimation of functional dependencies. The Hilbert envelope of each parcel’s time series was computed to extract the analytic signal, and epochs were concatenated. Pairwise functional connectivity between parcels was computed using amplitude envelope correlations (AEC), which was previously shown to be the most consistent network connectivity estimate at the group level (Colclough et al., 2016). The AEC of every pair of parcels formed the association matrices.

Choices of the analysis parameters were made based on test-retest reliability outcomes. The association matrices were thresholded at 25% density. This threshold was chosen because reliability of the metrics at low sparsities (< 10%) is low: networks get fractured and disconnected (Dennis et al., 2012). Reliability of the metrics is improved at higher densities (Braun et al., 2012), and have been shown to be stable between sparsities of 0.2 to 0.3, with a sharp drop in reliability above 0.3 (Dennis et al., 2012). We opted for thresholded weighted graphs as they generate more stable measurements compared to binarized graphs (Wang et al., 2011). Graph metrics were then calculated on the weighted association matrices using the Brain Connectivity Toolbox (Rubinov and Sporns, 2010) in Matlab 2017a (The Mathworks Inc., 2017). We use four metrics to capture essential global and local characteristics of the network communication at the nodal level. Metrics were computed for each of 98 nodes, five frequency bands and 12 participants, then normalized against 500 random graphs with equivalent degree.

(1) *Eigenvector centrality* is an extension of degree centrality. Degree centrality measures how many links connect with a node, giving equivalent weights to links coming from each connecting node. Eigenvector centrality is a meta-metric that quantifies the functional influence of a node on every other node in the graph, by weighting the importance of each nodal connection based on the influence of the nodes with which they connect. It is measured as the first eigenvector of the adjacency matrix corresponding to the largest eigenvalue (Bonacich, 1972).

(2) *Clustering coefficient* is the fraction of triangular connections formed by a node with other nodes. A node is strongly clustered if a large proportion of its neighbors are neighbors of each other. Since nodes that have high local clustering are also well connected locally, this measure captures local efficiency of information transfer.

(3) *Eccentricity* is defined as the longest distance between a node and any other node in the network. It is a nodal measure of global efficiency, reflecting long-range efficiency of information transfer and network integration, where high values of eccentricity indicate low global efficiency of communication.

(4) *Participation coefficient* reflects the diversity of nodes’ intermodular connections (i.e. connectivity to multiple functional modules), and is computed using the Louvain community detection algorithm (Blondel et al., 2008). Participation coefficient captures the segregation of functional networks, where a high participation coefficient would indicate connectivity to a high number of segregated functional modules. Modular networks maintain a balance between functionally specialized modules that have high within and between-module connectivity. Because of this fine balance, higher participation coefficient values does not necessarily correspond to better modularity; they could reflect a breakdown of functional segregation.

### 2.8. ROI selection

ROI selection was based on brain regions widely reported to accumulate neurofibrillary tangles and show atrophy in early stages of Alzheimer’s disease including Braak areas (Braak et al., 2006; Jack et al., 2018; Johnson et al., 2016; Ossenkoppele et al., 2016). ROIs consisted of hippocampus, entorhinal cortex, parahippocampal gyrus, angular gyrus, supramarginal gyrus, posterior and temporo-occipital middle and inferior temporal gyri, anterior and posterior fusiform cortex, precuneus, posterior cingulate gyrus, middle and superior frontal gyri and orbitofrontal cortex bilaterally.

### 2.9. Correlations and statistical analysis

We tested the relationship between the network properties and Tau burden both at the lobar level and at the level of the above ROIs. Metrics calculated for each participant were partially correlated with the Tau burden at each parcellation using Pearson’s correlations, whilst controlling for participants’ age. The nodes were grouped into five functional areas (i.e. lobes): frontal, temporal, parietal, occipital and limbic areas. To test changes at the lobe level, the correlation coefficients extracted from the nodes from each functional area were Fisher z-transformed then tested using one sample t-tests.

Finally, to explore longitudinal changes in network function, the differences in connectivity and graph metrics for the lobes and ROIs were compared between the baseline and 6-month visit using paired t-tests. In addition, to quantify the relationship between the longitudinal changes and Tau pathology, we correlated the Tau burden with these changes in connectivity. The resulting p values were corrected for multiple comparisons of metrics, bands and lobes, using false discovery rate at 0.05. The graph metrics in the MNI space were back-projected onto the canonical Freesurfer cortical surface for ease of visualization using the MNI2FS toolbox (https://github.com/dprice80/mni2fs).

## 3. Results

### 3.1. Tau deposition and connectivity

All participants showed the typical widespread bilateral Tau deposition (Passamonti et al., 2017; Schöll et al., 2016). Fig 2A shows [^18^F]AV1451 BP_ND_ maps across participants, with highest levels at the precuneus, posterior cingulate, posterior middle temporal, anterior fusiform, inferior parietal lobules, and the putamen. Fig 2B shows the [^18^F]AV1451 BP_ND_ binding for each participant, ordered from left to right in decreasing MMSE scores. Fig 2C displays the mean degree maps at the baseline visit, across the frequency bands. The maps were visualized using the BrainNet Viewer (www.nitrc.org/projects/bnv). There was strong connectivity between frontal, temporal and limbic nodes, and strong connectivity within occipital and parietal nodes.

**Fig 2.**
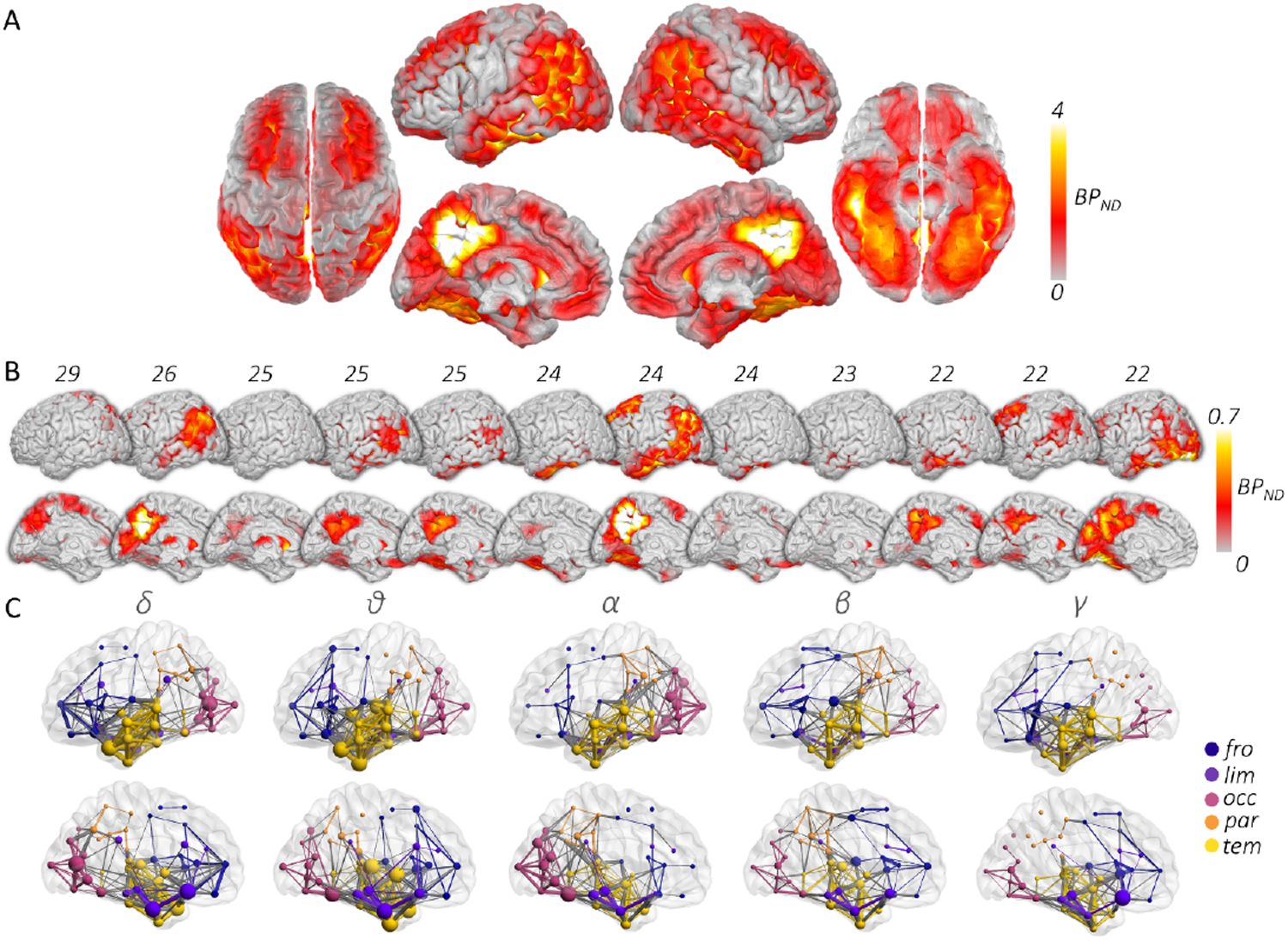
[18F] AV1451 binding maps and connectivity patterns. A. Map of tau deposition overlap across the participants, measured as the [^18^F]AV1451 non-displaceable binding potential where lighter colors indicate higher Tau burden across the sample. The map shows the characteristic AD tau distribution that spreads over temporoparietal, posterior medial and superior frontal areas bilaterally. Strongest overlap is observed around the precuneus, angular gyri, posterior middle temporal, and inferior temporal areas. B. Tau deposition maps of individual participants ordered by decreasing MMSE scores, displaying the left lateral and medial views. C. Degree maps displaying the mean connectivity patterns across five frequency bands on left lateral and medial surfaces. Connections are thresholded to display the strongest 10% for ease of visualization. Increasing node size and edge thickness indicate higher degree and stronger connectivity respectively. Note that the connectivity patterns across the bands largely overlap where the strongest connections are between the temporal nodes and the frontal and limbic nodes. Occipital nodes show strong within lobe, but weak between lobe connectivity. Parietal nodes show weak connectivity both within and between lobes.

### 3.2. Impact of Tau burden on network properties

Participation coefficient was significantly related to the Tau deposition in delta, theta, alpha and beta bands. Fig 3A shows the back-projected participation coefficient values at each node onto the cortical surface across five frequency bands. The values are thresholded for each frequency band to allow better visualization of cortical distribution. We find the highest values around the frontal, posterior temporal lobes as well as posterior cingulate gyrus and precuneus indicating that these areas have higher number of connections to functional modules in lower frequency bands. Fig 4A displays the distribution of correlation values between participation coefficient and Tau within functional lobes. The results of the lobar level correlations of the metrics with Tau are given in Table 1.

**Table 1.**
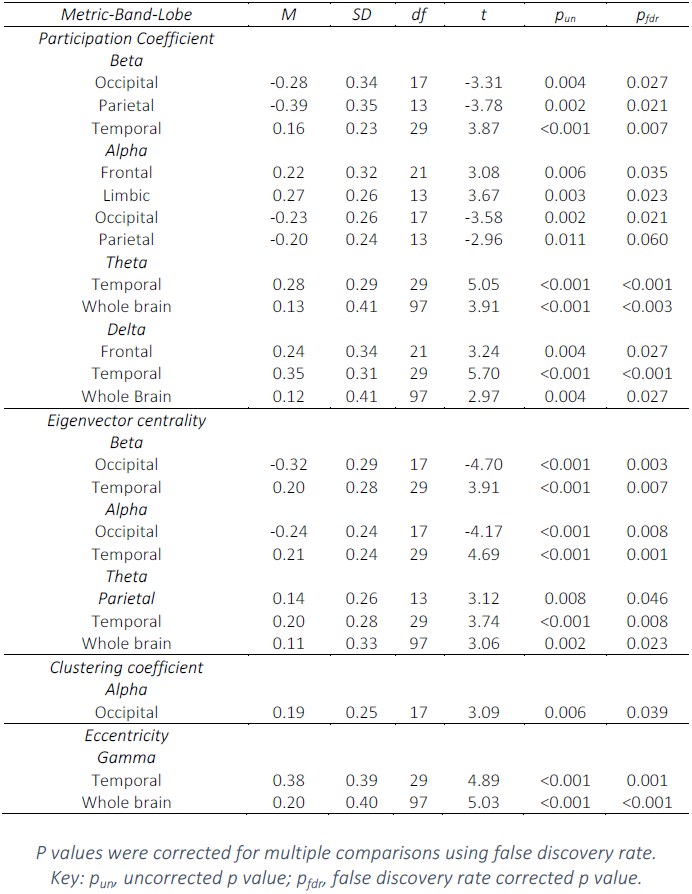
Lobar level analysis results across metrics and bands

| Metric-Band-Lobe | M | SD | df | t | $p_{un}$ | $p_{fdr}$ |
| --- | --- | --- | --- | --- | --- | --- |
| *Participation Coefficient* |  |  |  |  |  |  |
| *Beta* |  |  |  |  |  |  |
| Occipital | -0.28 | 0.34 | 17 | -3.31 | 0.004 | 0.027 |
| Parietal | -0.39 | 0.35 | 13 | -3.78 | 0.002 | 0.021 |
| Temporal | 0.16 | 0.23 | 29 | 3.87 | <0.001 | 0.007 |
| *Alpha* |  |  |  |  |  |  |
| Frontal | 0.22 | 0.32 | 21 | 3.08 | 0.006 | 0.035 |
| Limbic | 0.27 | 0.26 | 13 | 3.67 | 0.003 | 0.023 |
| Occipital | -0.23 | 0.26 | 17 | -3.58 | 0.002 | 0.021 |
| Parietal | -0.20 | 0.24 | 13 | -2.96 | 0.011 | 0.060 |
| *Theta* |  |  |  |  |  |  |
| Temporal | 0.28 | 0.29 | 29 | 5.05 | <0.001 | <0.001 |
| Whole brain | 0.13 | 0.41 | 97 | 3.91 | <0.001 | <0.003 |
| *Delta* |  |  |  |  |  |  |
| Frontal | 0.24 | 0.34 | 21 | 3.24 | 0.004 | 0.027 |
| Temporal | 0.35 | 0.31 | 29 | 5.70 | <0.001 | <0.001 |
| Whole Brain | 0.12 | 0.41 | 97 | 2.97 | 0.004 | 0.027 |
| *Eigenvector centrality* |  |  |  |  |  |  |
| *Beta* |  |  |  |  |  |  |
| Occipital | -0.32 | 0.29 | 17 | -4.70 | <0.001 | 0.003 |
| Temporal | 0.20 | 0.28 | 29 | 3.91 | <0.001 | 0.007 |
| *Alpha* |  |  |  |  |  |  |
| Occipital | -0.24 | 0.24 | 17 | -4.17 | <0.001 | 0.008 |
| Temporal | 0.21 | 0.24 | 29 | 4.69 | <0.001 | 0.001 |
| *Theta* |  |  |  |  |  |  |
| Parietal | 0.14 | 0.26 | 13 | 3.12 | 0.008 | 0.046 |
| Temporal | 0.20 | 0.28 | 29 | 3.74 | <0.001 | 0.008 |
| Whole brain | 0.11 | 0.33 | 97 | 3.06 | 0.002 | 0.023 |
| *Clustering coefficient* |  |  |  |  |  |  |
| *Alpha* |  |  |  |  |  |  |
| Occipital | 0.19 | 0.25 | 17 | 3.09 | 0.006 | 0.039 |
| *Eccentricity* |  |  |  |  |  |  |
| *Gamma* |  |  |  |  |  |  |
| Temporal | 0.38 | 0.39 | 29 | 4.89 | <0.001 | 0.001 |
| Whole brain | 0.20 | 0.40 | 97 | 5.03 | <0.001 | <0.001 |

*P* values were corrected for multiple comparisons using false discovery rate.
Key: $p_{un}$ , uncorrected *p* value; $p_{fdr}$ , false discovery rate corrected *p* value.

**Fig 3.**
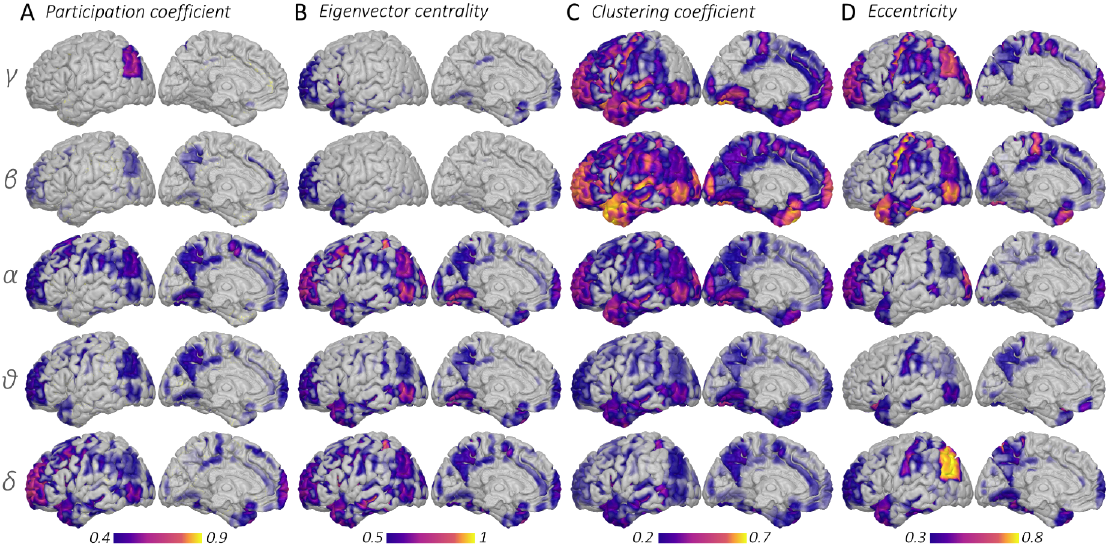
Distribution of normalized graph metrics across the cortex and frequency bands. 3D back-projections of the graph metrics onto the left lateral and medial cortical surfaces, going from gamma down to delta band. The scales are adjusted for each metric to allow better visualization of differences in cortical distribution. (A) Participation coefficient and (B) eigenvector centrality are higher for lower frequency bands. Their distribution is overlapping and concentrated around frontal, inferior parietal areas and precuneus. However, the efficiency measures (C-D), display higher values for higher frequency bands, suggesting that local and global efficiency are disrupted more strongly in lower and higher frequency bands respectively. Key: γ, gamma; β, beta; α, alpha; ϑ, theta; δ, delta band

**Fig 4.**
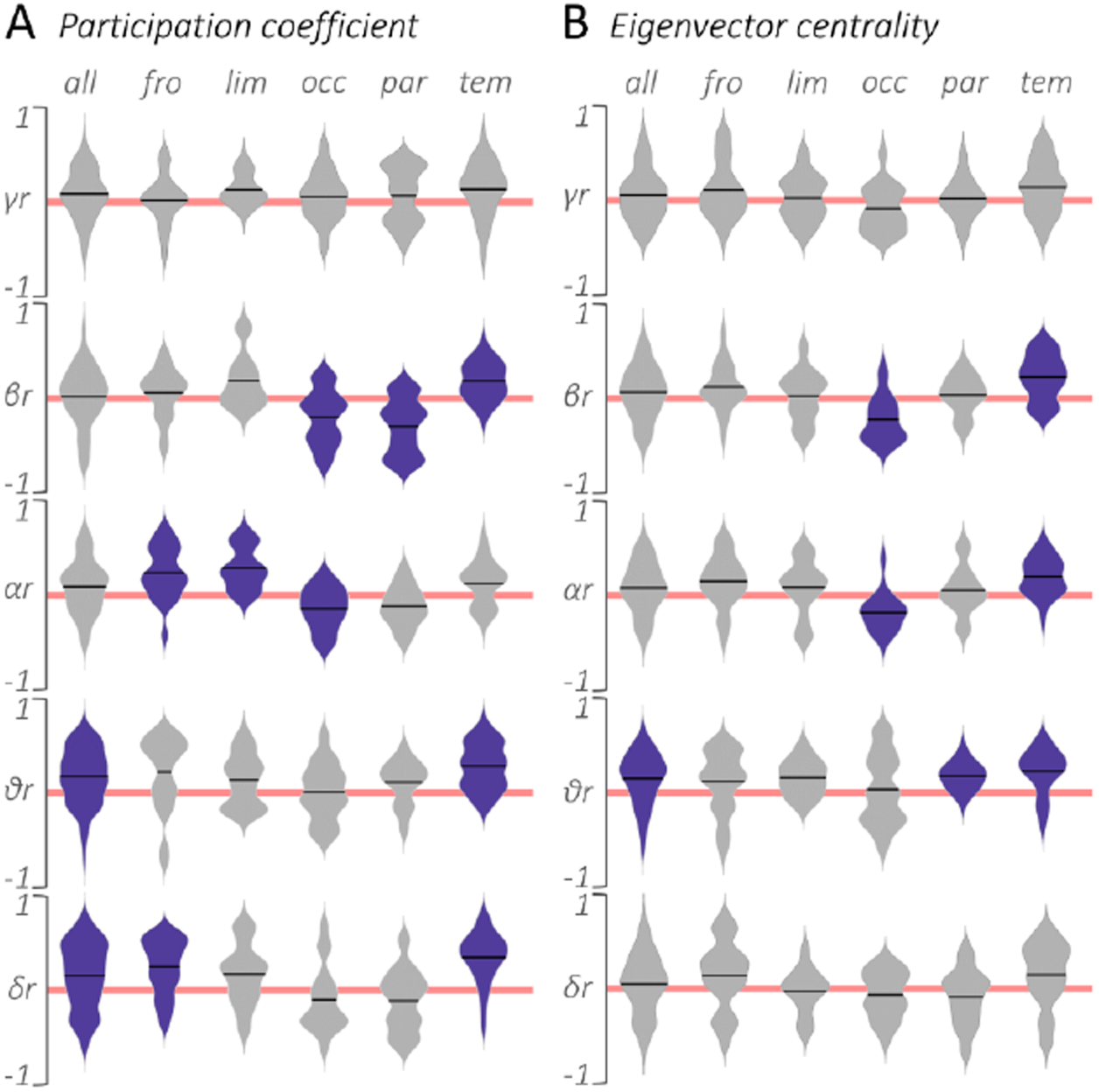
Lobar level correlations between metrics and Tau burden. Violin plots displaying the distribution of r values across parcels and frequency bands, clustered by functional lobes. Red line indicates r=0. Black lines indicate the mean. Correlations that were significantly different than zero after FDR correction are displayed in purple. A. Participation coefficient correlations. Note consistent positive correlations of tau with the frontal, limbic and temporal nodes, and negative correlations with the occipital and parietal nodes, indicating widespread changes in functional specificity and segregation. B. Eigenvector centrality correlations. Between theta and beta bands, the centrality of temporal and parietal nodes increases with increasing tau deposition, whereas occipital nodes are shown to have decreasing central role in communication. Key: all, whole brain; fro, frontal; lim, limbic; occ, occipital; par, parietal; tem, temporal; r, Pearson’s r; γ, gamma; β, beta; α, alpha; ϑ, theta; δ, delta band.

We found contrasting patterns between functional involvement and segregation with Tau across functional lobes. The participation coefficient in the frontal, limbic and temporal lobes showed a positive relationship with Tau burden, where this effect was stronger for lower frequency bands, suggesting increases in involvement of nodes in multiple functional modules, i.e. decreases in functional segregation of the cortex. We found the opposite pattern in the occipital and parietal nodes being stronger in higher frequency bands. Decreases in participation coefficient in these lobes indicate an increasing isolation from remaining functional networks in the brain. The ROI analysis results given in Table 2 corroborate this pattern and reveal that the strongest effects in participation coefficient among the Alzheimer’s ROIs are observed in the entorhinal and parahippocampal cortices.

**Table 2.** Regional analysis results

| Metric-Band-ROI | *r* | *p<sub>un</sub>* | *p<sub>fdr</sub>* |
| --- | --- | --- | --- |
| *Participation coefficient* |  |  |  |
| *Gamma* |  |  |  |
| L pos temporal fusiform gyrus | 0.75 | 0.008 | 0.031 |
| *Beta* |  |  |  |
| R entorhinal cortex | 0.85 | >0.001 | 0.003 |
| L parahippocampal cortex | 0.81 | 0.003 | 0.011 |
| R precuneus | -0.76 | 0.007 | 0.028 |
| *Alpha* |  |  |  |
| L orbitofrontal cortex | 0.76 | 0.007 | 0.027 |
| *Delta* |  |  |  |
| R pos temporal fusiform gyrus | 0.85 | 0.006 | 0.026 |
| *Eigenvector centrality* |  |  |  |
| *Gamma* |  |  |  |
| R superior frontal gyrus | 0.74 | 0.008 | 0.034 |
| R middle frontal gyrus | 0.80 | 0.003 | 0.013 |
| *Beta* |  |  |  |
| R entorhinal cortex | -0.69 | 0.017 | 0.035 |
| *Delta* |  |  |  |
| R pos supramarginal gyrus | -0.81 | 0.015 | 0.042 |
| *Clustering coefficient* |  |  |  |
| *Gamma* |  |  |  |
| L inferior temporo-occipital gyrus | 0.76 | 0.006 | 0.025 |
| R angular gyrus | -0.83 | 0.001 | 0.006 |
| R middle frontal gyrus | 0.75 | 0.008 | 0.016 |
| L superior frontal gyrus | 0.72 | 0.012 | 0.049 |
| *Beta* |  |  |  |
| R entorhinal cortex | -0.69 | 0.017 | 0.035 |
| *Alpha* |  |  |  |
| L parahippocampal cortex | -0.75 | 0.008 | 0.032 |
| *Delta* |  |  |  |
| R pos supramarginal gyrus | -0.78 | 0.021 | 0.042 |
| R inferior temporal gyrus | -0.83 | 0.011 | 0.046 |
| *Eccentricity* |  |  |  |
| *Gamma* |  |  |  |
| R pos inferior temporal gyrus | 0.81 | 0.007 | 0.031 |
| R pos temporal fusiform gyrus | 0.86 | 0.003 | 0.012 |
| R ant temporal fusiform gyrus | 0.79 | 0.010 | 0.042 |
| R orbitofrontal cortex | 0.79 | 0.011 | 0.045 |
| R entorhinal cortex | 0.77 | 0.014 | 0.056 |
| L pos inferior temporal gyrus | 0.72 | 0.028 | 0.056 |

*P* values were corrected for multiple comparisons using false discovery rate.
Key: $p_{un}$ , uncorrected *p* value; $p_{fdr}$ , false discovery rate corrected *p* value; R, right; L, left; pos, posterior; ant, anterior.

The eigenvector centrality captured a node’s central influence in communication between other central nodes. The values were higher for lower frequencies especially around bilateral middle and superior frontal as well as posterior temporal and cingulate and precuneus. We found significant effects (Table 1) in theta, alpha and beta bands (Fig 4B), with similar patterns of decreasing and increasing centrality in the occipital and temporal nodes respectively with increasing Tau levels. Regional analysis revealed (Table 2) positive correlations in the frontal nodes. However, in the absence of lobar level results, these effects indicate that the centrality changes in the frontal lobe are focal.

Clustering coefficient values displayed in Fig 3C were higher for high frequency bands, observed in the bilateral middle and superior frontal gyri, temporal poles and middle temporal gyri. In contrast to our predictions, correlations of clustering coefficient with Tau (see Supplementary information) were modest and restricted to the occipital nodes (Table 1). Regional analyses revealed a negative relationship in the temporal and limbic regions but a positive relationship in the frontal regions (Table 2). In the absence of strong lobar level effects, these results indicate that Tau-related clustering changes are more focal.

Finally, Fig 3D shows the back-projections of the eccentricity values on the cortex. We found a positive relationship between eccentricity and Tau in the temporal nodes solely in the gamma band, i.e. decreased global efficiency. Regional analyses further added negative correlations in the orbitofrontal and entorhinal cortex.

### 3.3. Longitudinal changes in network properties and connectivity

The connectivity changes within the lobes, connectivity of each ROI with the remaining Alzheimer’s ROIs (i.e. 31 nodes) displayed only subtle changes in magnitude and were not significant at the corrected level. At the lobar level only one effect survived the FDR correction. The frontal participation coefficient in the gamma band showed significant increases over 6 months (*t*(11) = −2.86; *p*_*un*_ = 0.009; *p*_*fdr*_ = 0.036).

ROI analyses showed significant results for eccentricity and eigenvector centrality metrics longitudinally. Eccentricity showed increases in 6 months period in the left posterior middle temporal gyrus in the gamma band (*t*(11) = −3.02; *p*_*un*_ = 0.007; *p*_*fdr*_ = 0.035), providing evidence for further decreases in long range efficiency of gamma synchrony. We found significant decreases in node centrality as measured by the eigenvector centrality in left precuneus (*t*(11) = 3.57; *p*_*un*_ = 0.003; *p*_*fdr*_ = 0.013) and posterior middle temporal gyrus (*t*(11) = 3.18; *p*_*un*_ = 0.006; *p*_*fdr*_ = 0.029) and a marginal effect in the left angular gyrus (*t*(11) = 2.83; *p*_*un*_ = 0.012; *p*_*fdr*_ = 0.060) all in delta band. These eigenvector centrality results complement the negative correlations shown for eigenvector centrality-Tau in the parietal nodes.

### 3.4. Tau-related rates of change in networks

We further tested the relationship between the connectivity and graph metric changes over time and the baseline Tau burden. Negative and positive correlations indicate overall patterns of increases and decreases in measurements over a 6 months’ period in relation to the participants’ Tau levels at the baseline visit. The change in delta connectivity between temporal and frontal (*r*_*Δ*_ = −0.85; *p*_*un*_ = 0.016; *p*_*fdr*_ = 0.018), limbic (*r* = −0.85; *p*_*un*_ = 0.015; *p*_*fdr*_ = 0.018), occipital (*r*_*Δ*_ = −0.84; *p*_*un*_ = 0.017; *p*_*fdr*_ = 0.018), and parietal (*r*_*Δ*_ = −0.85; *p*_*un*_ = 0.015; *p*_*fdr*_ =0.018) lobes, as well as within the temporal lobe itself (*r*_*Δ*_ = −0.84; *p*_*un*_ = 0.018; *p*_*fdr*_ = 0.018), significantly correlated with the mean Tau burden in the temporal lobe. We then focused on the correlations at the ROI level. The delta band connectivity changes of the right entorhinal cortex (*r*_*Δ*_ = −0.94; *p*_*un*_ = 0.002; *p*_*fdr*_ = 0.008), posterior temporal fusiform cortex (*r*_*Δ*_ = −0.90; *p*_*un*_ = 0.005; *p*_*fdr*_ = 0.025), and posterior inferior temporal gyrus (*r*_*Δ*_ = −0.91; *p*_*un*_ = 0.004; *p*_*fdr*_ = 0.021) with the remaining ROIs was significantly related to their Tau burden. Further, in the theta band the right temporo-occipital inferior temporal gyrus showed significant correlations (*r*_*Δ*_ = −0.85; *p*_*un*_ = 0.007; *p*_*fdr*_ = 0.036).

We did not find any significant correlations of the graph metrics at the lobe level, indicating that Tau related rate of change in metrics is not equally distributed across the functional lobes. However, we found significant correlations among the ROIs. The change in participation coefficient related to the Tau burden in the right superior frontal gyrus (*r*_*Δ*_ = 0.91; *p*_*un*_ = 0.004; *p*_*fdr*_ = 0.044) in the delta band, suggesting a decrease in functional involvement. Eigenvector centrality showed effects in the delta and beta bands. In the delta band left middle frontal gyrus (*r*_*Δ*_ = 0.93; *p*_*un*_ = 0.002; *p*_*fdr*_ = 0.045) and in the beta band the left posterior temporal fusiform cortex (*r*_*Δ*_ = 0.89; *p*_*un*_ = 0.001; *p*_*fdr*_ = 0.025) showed a significant effect, indicating decreases in their centrality in information transfer between important nodes. Clustering coefficient rate of change significantly related to Tau in the right superior frontal gyrus (*r*_*Δ*_ = −0.91; *p*_*un*_ = 0.004; *p*_*fdr*_ = 0.044) in the delta band, suggesting an increase in the local efficiency of information transfer to the baseline Tau burden. We did not find significant relationship between rate of change in eccentricity and the baseline Tau.

## 4. Discussion

In this study, we investigated the impact of Tau burden on neurophysiological network properties as captured by E/MEG, and the effectiveness of these biomarkers to track short-term disease progression in early Alzheimer’s disease. We demonstrated that the segregation of functional brain networks and the functional influence of regions exerted on the remaining network were modulated as a function of Tau burden, leading to an increasingly fragmented network. Secondly, we showed that Tau burden disrupts global more than local efficiency of information transfer. We finally report key regions that display greatest shifts in network properties in 6 months and in relation to the Tau burden: including middle temporal and angular gyri, superior frontal gyri and precuneus. Our findings provide support for neurophysiological biomarkers in experimental medicines studies or early phase trials in Alzheimer’s disease.

In line with our predictions, the analysis showed Tau-related increases in participation coefficient in the frontal, limbic and temporal lobes at lower frequencies (delta and theta bands). This is in agreement with previous studies focusing on the mild cognitive impairment and Alzheimer’s disease patients that report increases in functional modules in the delta and theta bands (de Haan et al., 2012a), and increased global participation in fMRI (Cope et al., 2018). Increases in participation coefficient suggest an increased involvement in functional modules. A healthy profile of functional involvement, hinges on a fine balance between inter- and intra-modular communication, where the breakdown of this profile could be a result of either a sub-optimal increase (i.e. disruption of segregation), or decrease (i.e. isolation) of functional involvement. Thus, widespread increases in participation coefficient suggest disruptions in functional segregation and sub-optimal network processing. We found the opposite pattern in the parietal and occipital lobes, which indicates a disruption of their multi-module connectivity and increasing isolation as the pathology progressed. This interpretation is in line with the findings of a large scale MR study by Brier and colleagues (2014) which showed decreasing participation coefficient and inter-modular connectivity in the frontal, occipital and parietal regions in mild cognitive impairment and Alzheimer’s disease patients compared to controls. Similarly, in a hidden markov modeling analysis, the posterior default mode network of Alzheimer’s patients, consisting of precuneus and posterior cingulate cortex, was visited less often and for shorter periods (Sitnikova et al., 2018).

The eigenvector centrality results complemented the participation coefficient results showing Tau-related decreases of the occipital nodes and increases of the temporal and frontal nodes. This fronto-occipital pattern has been recently reported using fMRI, related to the CSF p-Tau level as well as patients’ MMSE scores (Binnewijzend et al., 2014; Cope et al., 2018). Similarly, the glucose metabolism in the occipital areas of the ApoE4 carriers gets reduced (Ossenkoppele et al., 2013). Synchronization likelihood of the occipital areas decrease whereas a increase is observed among the frontal nodes (Sanz-Arigita et al., 2010). Complementary to what we found for participation coefficient, these results suggest a shift towards an increasingly fragmented network where frontal and temporal nodes become more influential and involved in information transfer, whereas parietal and occipital nodes get further disconnected and isolated.

Eccentricity showed strong disruptions among the temporal nodes in the highest frequencies (gamma band), suggesting an impairment in global efficiency of feed-forward information transfer. These findings are consistent with previous reports showing increases in the global characteristic path length (Zhao et al., 2012) with decreasing MMSE scores (Stam et al., 2006), and decreases in global efficiency and connectivity between long-distance hubs (Li et al., 2013; Liu et al., 2014). Long range gamma synchrony is thought to be a fundamental function of the brain serving to integrate information processed in-tandem across regions in a network (Başar-Eroglu et al., 1996). Synchronization of processing in the gamma band exists both locally and across long distances in the cortex even with zero lag delays (Rodriguez et al., 1999; Singer, 1999). Previous studies reported widespread loss of long-range gamma synchrony in humans (Koenig et al., 2005; Stam et al., 2002) and tau mediated network instability in gamma band in mouse models of Alzheimer’s disease (Verret et al., 2012). We speculate that the effect in the gamma band results from the degeneration of cholinergic projections and impaired muscarinic reception function, which enhance gamma frequencies in the healthy brain (Rodriguez et al., 2004).

The secondary aim of the study was to assess and quantify changes that occur over 6 months. Changes in such a short period could aid evaluation and fast turnover of experimental drugs for Alzheimer’s disease. Among our four network metrics, all but clustering coefficient showed significant changes in 6 months, where we observed largest changes for the eigenvector centrality. Eigenvector centrality showed decreases in the precuneus, middle temporal gyrus and marginally in the angular gyrus in the delta band, indicating decreases of functional influence over other nodes in the network. In line with the lobar level findings, we found further decreases in global efficiency, in the posterior middle temporal gyrus in gamma band. Finally, in contrast to the widespread effects at the lobar level, participation coefficient displayed modest increases in the gamma band only in the frontal lobe. Among the four metrics, only two (i.e. eigenvector centrality and eccentricity) showed direct relationship to the local tau burden, whilst capturing short-term changes in network functioning. This analysis also highlighted key regions that show faster rates of change in their network properties over 6 months. We found that the network properties changed faster in the precuneus, inferior parietal lobule, and the middle temporal gyrus compared to the rest of the network. This could be attributed to their faster rates of Tau accumulation (Ishiki et al., 2015) and of cortical thinning observed for the mild cognitive impairment patients converting to Alzheimer’s disease and Alzheimer’s patients (Li et al., 2012).

The rate of change of connectivity related to Tau burden at baseline. The temporal Tau levels at baseline were linked to increased delta connectivity of the temporal lobe with the remaining network as well as within itself. Similarly, at the ROI level, we found increases in delta and theta connectivity of the entorhinal, inferior temporal and fusiform areas with the remaining network. These results are complementary to reported increases in delta and theta synchrony (Babiloni et al., 2004; Poza et al., 2008), where the slowing of frequencies was related to the white matter atrophy (Babiloni et al., 2006) and progression from mild cognitive impairment to Alzheimer’s disease (Babiloni et al., 2010; Huang et al., 2000; Jelic et al., 2000). Compared to Alzheimer’s patients who are either ε2 or ε3 carriers, ε4 carriers display longitudinal increases in delta and theta power (Lehtovirta et al., 1996). This slowing could be linked to the impairments in cholinergic-muscarinic transmission which causes decreases in gamma, and increases in resting delta and theta power (Bosboom et al., 2009).

We predicted that two measures of efficiency, clustering coefficient and eccentricity would display widespread disruptions in relation to Tau accumulation. In contrast to our predictions, we found focal effects for the clustering coefficient. The reports on local efficiency changes in Alzheimer’s disease measured by the clustering coefficient are mixed. Some reported decreases pronounced in the hippocampi (Supekar et al., 2008), and in the alpha and beta bands (de Haan et al., 2009), which relate to the positive biomarker status of the patients (Brier et al., 2014). Others showed increases in the delta and alpha bands (Buldú et al., 2011; Vecchio et al., 2014; Zhao et al., 2012), or reported that the clustering remains relatively spared in disease progression (Cope et al., 2018; Stam et al., 2006). Our regional analyses showed a Tau related negative pattern in medial temporal and inferior parietal areas, and a positive pattern in the middle frontal and temporo-occipital areas. The areas showing a reduction in clustering were entorhinal, parahippocampal cortex and angular and supramarginal gyri which accumulate Tau earlier in disease. We attribute this pattern to a non-linear impact of tau on local clustering; i.e. as Tau builds up, the clustering initially increases followed by a decrease, which could additionally explain the abundant contradicting findings in the literature.

E/MEG has been widely utilized in network research. Its use historically originates from the field of epilepsy, however now it is used for in-depth understanding of neurophysiological underpinnings of the fundamental brain processes and their dysfunction. Despite its lower spatial resolution compared to MR and PET, E/MEG remains the only scanner that can directly measure activity produced by neuronal populations, and it does so at the brain’s inherent speed, i.e. the millisecond time resolution. This fast capture allows us to see transient patterns of brain activity invisible to slower scanners such as MR. Moreover, it allows investigations of brain activity patterns, spread across different frequency bands. The current analysis similarly demonstrates this multi-faceted nature of neurophysiology, i.e. how connectivity patterns and network metrics behave differently to low versus high frequency bands. The information embedded in different frequencies could be valuable in detecting disorders, where neuronal populations show impaired functions such as in neurodegenerative disorders.

When EEG and MEG are combined, they could capture the brain activity originating from both radial and tangential orientations brain tissue. Since the sensors are outside the head, the signal to noise ratio decreases as the distance between the sensors and the electrical source in the brain increases. This makes the signals estimated from subcortical nuclei and ventral areas weaker compared to more lateral and superficial sources. However, the current results indicate that despite having lower signal to noise and weaker amplitudes, we are able to capture patterns of brain activity in deeper sources even with a small sample of participants.

Due to the small sample size of the pilot study and the absence of data from healthy controls, we have adopted an exploratory correlational approach between the Tau burden and neurophysiological metrics. However, future confirmatory studies could increase the number of patients and include data from matched healthy participants. Similarly, acquiring Tau PET at two time points, and relating changes in longitudinal Tau PET to the changes in longitudinal graph metrics, would strengthen the findings.

## 5. Conclusions

Our findings provide the first evidence that Tau pathology disrupts human brain network connectivity. This may be due to synaptic dysfunction as shown in animal models, or additionally from somatic and axonal deficits. In early Alzheimer’s disease with increasing Tau burden, the brain shifts towards a fragmented network where fronto-temporal areas became crucial for information transfer, and parietal and occipital areas get further disconnected from the remaining network. Whilst we observe modest local disruptions in efficiency, Alzheimer’s disease displays a greater impact on long-range global efficiency of the temporal areas. Since neurophysiological network biomarkers directly relate to Tau pathology, they could be utilized as a non-invasive tool to track short term disease progression and the impact of disease modifying therapies.

## Abbreviations

Aβ: Amyloid-beta;
AEC: Amplitude envelope correlations;
BP_ND_: Non-displaceable binding potential;
CSF: Cerebrospinal fluid;
EEG: Electroencephalography;
FBP: Filtered back projection;
FSL: FMRIB software library;
HO: Harvard-Oxford Atlas;
ICA: Independent component analysis;
MEG: Magnetoencephalography;
MMSE: Mini mental state examination;
MRI: Magnetic resonance imaging;
NFT: Neurofibrillary tangles;
NINCDS/ADRDA: National Institute of Neurological and Communicative Disorders and Stroke and the Alzheimer’s Disease and Related Disorders Association;
PET: Positron emission tomography;
PHF: Paired helical filaments;
P-tau: Hyper-phosphorylated tau;
SPM: Statistical parametric mapping;
SRTM: Simplified reference tissue model;
TAC: Time activity curve

## Author’s contributions

S.L., J.B.R, R.N.H., G.G., G.B., M.W.W., and R.G. designed the experiment. A.Q., E.C., A.G. collected the E/MEG data. A.F. pre-processed the PET and MR data. E.K. pre-processed the MEG data, performed the data analysis. E.K. and J.B.R. wrote the manuscript, and all authors contributed to the final version.

## Acknowledgments

The Deep and Frequent Phenotyping Study is funded by the Medical Research Council and National Institute for Health Research as part of the Dementias Platform UK (MR/N029941/1). JBR is supported by the Wellcome Trust (103838) and Medical Research Council (SUAG/004 RG91365). RH is supported by the Medical Research Council (SUAG/010 RG91365). MWW’s research is supported by the NIHR Oxford Health Biomedical Research Centre, the Wellcome Trust (106183/Z/14/Z and 203139/Z/16/Z) and the MRC UK MEG Partnership Grant (MR/K005464/1). EK is funded by the Dementias Platform UK and Alzheimer’s Research UK (RG94383/RG89702). We thank all participants and their families, the PET technicians, radiochemists, the MRI radiographers, and the clinical research nurses for their cooperation and support of this study. We thank Avid radiopharmaceuticals for the provision of [^18^F]AV1451 doses, and Wellcome Centre for Human Neuroimaging at University College London for their support with MEG scans. We thank Dr Rezvan Farahibozorg for useful discussions about the analyses and methods.

## References

Ahmed, Z., Cooper, J., Murray, T.K., Garn, K., McNaughton, E., Clarke, H., Parhizkar, S., Ward, M.A., Cavallini, A., Jackson, S., Bose, S., Clavaguera, F., Tolnay, M., Lavenir, I., Goedert, M., Hutton, M.L., O’Neill, M.J., 2014. A novel in vivo model of tau propagation with rapid and progressive neurofibrillary tangle pathology: the pattern of spread is determined by connectivity, not proximity. Acta neuropathologica 127(5), 667–683.

Babiloni, C., Binetti, G., Cassetta, E., Cerboneschi, D., Dal Forno, G., Del Percio, C., Ferreri, F., Ferri, R., Lanuzza, B., Miniussi, C., Moretti, D.V., Nobili, F., Pascual-Marqui, R.D., Rodriguez, G., Romani, G.L., Salinari, S., Tecchio, F., Vitali, P., Zanetti, O., Zappasodi, F., Rossini, P.M., 2004. Mapping distributed sources of cortical rhythms in mild Alzheimer’s disease. A multicentric EEG study. Neuroimage 22(1), 57–67.

Babiloni, C., Frisoni, G., Steriade, M., Bresciani, L., Binetti, G., Del Percio, C., Geroldi, C., Miniussi, C., Nobili, F., Rodriguez, G., Zappasodi, F., Carfagna, T., Rossini, P.M., 2006. Frontal white matter volume and delta EEG sources negatively correlate in awake subjects with mild cognitive impairment and Alzheimer’s disease. Clin Neurophysiol 117(5), 1113–1129.

Babiloni, C., Frisoni, G.B., Vecchio, F., Pievani, M., Geroldi, C., De Carli, C., Ferri, R., Vernieri, F., Lizio, R., Rossini, P.M., 2010. Global functional coupling of resting EEG rhythms is related to white-matter lesions along the cholinergic tracts in subjects with amnesic mild cognitive impairment. J Alzheimers Dis 19(3), 859–871.

Başar-Eroglu, C., Strüber, D., Schürmann, M., Stadler, M., Başar, E., 1996. Gamma-band responses in the brain: a short review of psychophysiological correlates and functional significance. Int J Psychophysiol 24(1-2), 101–112.

Binnewijzend, M.A., Adriaanse, S.M., Van der Flier, W.M., Teunissen, C.E., de Munck, J.C., Stam, C.J., Scheltens, P., van Berckel, B.N., Barkhof, F., Wink, A.M., 2014. Brain network alterations in Alzheimer’s disease measured by eigenvector centrality in fMRI are related to cognition and CSF biomarkers. Hum Brain Mapp 35(5), 2383–2393.

Blondel, V.D., Guillaume, J.-L., Lambiotte, R., Lefebvre, E., 2008. Fast unfolding of communities in large networks. Journal of Statistical Mechanics: Theory and Experiment P10008, 1–12.

Bonacich, P., 1972. Factoring and weighting approaches to clique identification. J Math Soc 2, 113–120.

Bosboom, J.L., Stoffers, D., Stam, C.J., Berendse, H.W., Wolters, E.C.h., 2009. Cholinergic modulation of MEG resting-state oscillatory activity in Parkinson’s disease related dementia. Clin Neurophysiol 120(5), 910–915.

Braak, H., Alafuzoff, I., Arzberger, T., Kretzschmar, H., Del Tredici, K., 2006. Staging of Alzheimer disease-associated neurofibrillary pathology using paraffin sections and immunocytochemistry. Acta Neuropathol 112(4), 389–404.

Braun, U., Plichta, M.M., Esslinger, C., Sauer, C., Haddad, L., Grimm, O., Mier, D., Mohnke, S., Heinz, A., Erk, S., Walter, H., Seiferth, N., Kirsch, P., Meyer-Lindenberg, A., 2012. Test-retest reliability of resting-state connectivity network characteristics using fMRI and graph theoretical measures. Neuroimage 59(2), 1404–1412.

Brier, M.R., Gordon, B., Friedrichsen, K., McCarthy, J., Stern, A., Christensen, J., Owen, C., Aldea, P., Su, Y., Hassenstab, J., Cairns, N.J., Holtzman, D.M., Fagan, A.M., Morris, J.C., Benzinger, T.L., Ances, B.M., 2016. Tau and Aβ imaging, CSF measures, and cognition in Alzheimer’s disease. Sci Transl Med 8(338), 338ra366.

Brier, M.R., Thomas, J.B., Fagan, A.M., Hassenstab, J., Holtzman, D.M., Benzinger, T.L., Morris, J.C., Ances, B.M., 2014. Functional connectivity and graph theory in preclinical Alzheimer’s disease. Neurobiol Aging 35(4), 757–768.

Buldú, J.M., Bajo, R., Maestú, F., Castellanos, N., Leyva, I., Gil, P., Sendiña-Nadal, I., Almendral, J.A., Nevado, A., del-Pozo, F., Boccaletti, S., 2011. Reorganization of functional networks in mild cognitive impairment. PLoS One 6(5), e19584.

Bullmore, E., Sporns, O., 2009. Complex brain networks: graph theoretical analysis of structural and functional systems. Nat Rev Neurosci 10(3), 186–198.

Colclough, G.L., Brookes, M.J., Smith, S.M., Woolrich, M.W., 2015. A symmetric multivariate leakage correction for MEG connectomes. Neuroimage 117, 439–448.

Colclough, G.L., Woolrich, M.W., Tewarie, P.K., Brookes, M.J., Quinn, A.J., Smith, S.M., 2016. How reliable are MEG resting-state connectivity metrics? NeuroImage 138, 284–293.

Cope, T.E., Rittman, T., Borchert, R.J., Jones, P.S., Vatansever, D., Allinson, K., Passamonti, L., Vazquez Rodriguez, P., Bevan-Jones, W.R., O’Brien, J.T., Rowe, J.B., 2018. Tau burden and the functional connectome in Alzheimer’s disease and progressive supranuclear palsy. Brain 141(2), 550–567.

Dauwels, J., Vialatte, F., Musha, T., Cichocki, A., 2010. A comparative study of synchrony measures for the early diagnosis of Alzheimer’s disease based on EEG. Neuroimage 49(1), 668–693.

de Haan, W., Pijnenburg, Y.A., Strijers, R.L., van der Made, Y., van der Flier, W.M., Scheltens, P., Stam, C.J., 2009. Functional neural network analysis in frontotemporal dementia and Alzheimer’s disease using EEG and graph theory. BMC Neurosci 10, 101.

de Haan, W., van der Flier, W.M., Koene, T., Smits, L.L., Scheltens, P., Stam, C.J., 2012a. Disrupted modular brain dynamics reflect cognitive dysfunction in Alzheimer’s disease. Neuroimage 59(4), 3085–3093.

de Haan, W., van der Flier, W.M., Wang, H., Van Mieghem, P.F., Scheltens, P., Stam, C.J., 2012b. Disruption of functional brain networks in Alzheimer’s disease: what can we learn from graph spectral analysis of resting-state magnetoencephalography? Brain Connect 2(2), 45–55.

Delorme, A., Makeig, S., 2004. EEGLAB: an open source toolbox for analysis of single-trial EEG dynamics including independent component analysis. J Neurosci Methods 134(1), 9–21.

Dennis, E.L., Jahanshad, N., Toga, A.W., McMahon, K.L., De Zubicaray, G.I., Martin, N.G., Wright, M.J., Thompson, P.M., 2012. Test-retest reliability of graph theory measures of structural brain connectivity., International Conference on Medical Image Computing and Computer-Assisted Intervention. Springer, pp. 305–312.

Firouzian, A., Whittington, A., Searle, G.E., Koychev, I., Zamboni, G., Lovestone, S., 2018. Imaging Aβ and tau in early stage Alzheimer’s disease with [18F]AV45 and [18F]AV1451. EJNMMI Research 8:19.

Gonzalez-Escamilla, G., Lange, C., Teipel, S., Buchert, R., Grothe, M.J., Initiative, A.s.D.N., 2017. PETPVE12: an SPM toolbox for Partial Volume Effects correction in brain PET - Application to amyloid imaging with AV45-PET. Neuroimage 147, 669–677.

Henson, R.N., Mouchlianitis, E., Friston, K.J., 2009. MEG and EEG data fusion: simultaneous localisation of face-evoked responses. Neuroimage 47(2), 581–589.

Hsieh, H., Boehm, J., Sato, C., Iwatsubo, T., Tomita, T., Sisodia, S., Malinow, R., 2006. AMPAR removal underlies Abeta-induced synaptic depression and dendritic spine loss. Neuron 52(5), 831–843.

Huang, C., Wahlund, L., Dierks, T., Julin, P., Winblad, B., Jelic, V., 2000. Discrimination of Alzheimer’s disease and mild cognitive impairment by equivalent EEG sources: a cross-sectional and longitudinal study. Clin Neurophysiol 111(11), 1961–1967.

Ishiki, A., Okamura, N., Furukawa, K., Furumoto, S., Harada, R., Tomita, N., Hiraoka, K., Watanuki, S., Ishikawa, Y., Tago, T., Funaki, Y., Iwata, R., Tashiro, M., Yanai, K., Kudo, Y., Arai, H., 2015. Longitudinal Assessment of Tau Pathology in Patients with Alzheimer’s Disease Using [18F]THK-5117 Positron Emission Tomography. PLoS One 10(10), e0140311.

Ittner, L.M., Ke, Y.D., Delerue, F., Bi, M., Gladbach, A., van Eersel, J., Wolfing, H., Chieng, B.C., Christie, M.J., Napier, I.A., Eckert, A., Staufenbiel, M., Hardeman, E., Gotz, J., 2010. Dendritic function of tau mediates amyloid-beta toxicity in Alzheimer’s disease mouse models. Cell 142(3), 387–397.

Jack, C.R., Wiste, H.J., Schwarz, C.G., Lowe, V.J., Senjem, M.L., Vemuri, P., Weigand, S.D., Therneau, T.M., Knopman, D.S., Gunter, J.L., Jones, D.T., Graff-Radford, J., Kantarci, K., Roberts, R.O., Mielke, M.M., Machulda, M.M., Petersen, R.C., 2018. Longitudinal tau PET in ageing and Alzheimer’s disease. Brain 141(5), 1517–1528.

Jelic, V., Johansson, S.E., Almkvist, O., Shigeta, M., Julin, P., Nordberg, A., Winblad, B., Wahlund, L.O., 2000. Quantitative electroencephalography in mild cognitive impairment: longitudinal changes and possible prediction of Alzheimer’s disease. Neurobiol Aging 21(4), 533–540.

Jenkinson, M., Beckmann, C.F., Behrens, T.E., Woolrich, M.W., Smith, S.M., 2012. FSL. Neuroimage 62(2), 782–790.

Johnson, K.A., Schultz, A., Betensky, R.A., Becker, J.A., Sepulcre, J., Rentz, D., Mormino, E., Chhatwal, J., Amariglio, R., Papp, K., Marshall, G., Albers, M., Mauro, S., Pepin, L., Alverio, J., Judge, K., Philiossaint, M., Shoup, T., Yokell, D., Dickerson, B., Gomez-Isla, T., Hyman, B., Vasdev, N., Sperling, R., 2016. Tau positron emission tomographic imaging in aging and early Alzheimer disease. Ann Neurol 79(1), 110–119.

Kimura, T., Whitcomb, D.J., Jo, J., Regan, P., Piers, T., Heo, S., Brown, C., Hashikawa, T., Murayama, M., Seok, H., Sotiropoulos, I., Kim, E., Collingridge, G.L., Takashima, A., Cho, K., 2014. Microtubule-associated protein tau is essential for long-term depression in the hippocampus. Philosophical transactions of the Royal Society of London. Series B, Biological sciences 369(1633), 20130144.

Koenig, T., Prichep, L., Dierks, T., Hubl, D., Wahlund, L.O., John, E.R., Jelic, V., 2005. Decreased EEG synchronization in Alzheimer’s disease and mild cognitive impairment. Neurobiol Aging 26(2), 165–171.

Koss, D.J., Robinson, L., Drever, B.D., Plucińska, K., Stoppelkamp, S., Veselcic, P., Riedel, G., Platt, B., 2016. Mutant Tau knock-in mice display frontotemporal dementia relevant behaviour and histopathology. Neurobiol Dis 91, 105–123.

Koychev, I., Gunn, R.N., Firouzian, A., Lawson, J., Zamboni, G., Ridha, B., Sahakian, B.J., Rowe, J.B., Thomas, A., Rochester, L., Ffytche, D., Howard, R., Zetterberg, H., MacKay, C., Lovestone, S., (, D.a.F.P.s.t., 2017. PET Tau and Amyloid-β Burden in Mild Alzheimer’s Disease: Divergent Relationship with Age, Cognition, and Cerebrospinal Fluid Biomarkers. J Alzheimers Dis 60(1), 283–293.

Kurudenkandy, F.R., Zilberter, M., Biverstål, H., Presto, J., Honcharenko, D., Strömberg, R., Johansson, J., Winblad, B., Fisahn, A., 2014. Amyloid-β-induced action potential desynchronization and degradation of hippocampal gamma oscillations is prevented by interference with peptide conformation change and aggregation. J Neurosci 34(34), 11416–11425.

LaFerla, F.M., Oddo, S., 2005. Alzheimer’s disease: Abeta, tau and synaptic dysfunction. Trends Mol Med 11(4), 170–176.

Lehtovirta, M., Partanen, J., Könönen, M., Soininen, H., Helisalmi, S., Mannermaa, A., Ryynänen, M., Hartikainen, P., Riekkinen, P., 1996. Spectral analysis of EEG in Alzheimer’s disease: relation to apolipoprotein E polymorphism. Neurobiol Aging 17(4), 523–526.

Li, G., Bien-Ly, N., Andrews-Zwilling, Y., Xu, Q., Bernardo, A., Ring, K., Halabisky, B., Deng, C., Mahley, R.W., Huang, Y., 2009. GABAergic interneuron dysfunction impairs hippocampal neurogenesis in adult apolipoprotein E4 knockin mice. Cell Stem Cell 5(6), 634–645.

Li, S., Hong, S., Shepardson, N.E., Walsh, D.M., Shankar, G.M., Selkoe, D., 2009. Soluble oligomers of amyloid Beta protein facilitate hippocampal long-term depression by disrupting neuronal glutamate uptake. Neuron 62(6), 788–801.

Li, Y., Qin, Y., Chen, X., Li, W., 2013. Exploring the functional brain network of Alzheimer’s disease: based on the computational experiment. PLoS One 8(9), e73186.

Li, Y., Wang, Y., Wu, G., Shi, F., Zhou, L., Lin, W., Shen, D., Initiative, A.s.D.N., 2012. Discriminant analysis of longitudinal cortical thickness changes in Alzheimer’s disease using dynamic and network features. Neurobiol Aging 33(2), 427.e415-430.

Liu, L., Wong, T.P., Pozza, M.F., Lingenhoehl, K., Wang, Y., Sheng, M., Auberson, Y.P., Wang, Y.T., 2004. Role of NMDA receptor subtypes in governing the direction of hippocampal synaptic plasticity. Science 304(5673), 1021–1024.

Liu, Y., Yu, C., Zhang, X., Liu, J., Duan, Y., Alexander-Bloch, A.F., Liu, B., Jiang, T., Bullmore, E., 2014. Impaired long distance functional connectivity and weighted network architecture in Alzheimer’s disease. Cereb Cortex 24(6), 1422–1435.

McKhann, G., Drachman, D., Folstein, M., Katzman, R., Price, D., Stadlan, E.M., 1984. Clinical diagnosis of Alzheimer’s disease: report of the NINCDS-ADRDA Work Group under the auspices of Department of Health and Human Services Task Force on Alzheimer’s Disease. Neurology 34(7), 939–944.

Murray, M.E., Lowe, V.J., Graff-Radford, N.R., Liesinger, A.M., Cannon, A., Przybelski, S.A., Rawal, B., Parisi, J.E., Petersen, R.C., Kantarci, K., Ross, O.A., Duara, R., Knopman, D.S., Jack, C.R., Jr., Dickson, D.W., 2015. Clinicopathologic and 11C-Pittsburgh compound B implications of Thal amyloid phase across the Alzheimer’s disease spectrum. Brain : a journal of neurology 138(Pt 5), 1370–1381.

Ochoa, J.F., Alonso, J.F., Duque, J.E., Tobón, C.A., Mañanas, M.A., Lopera, F., Hernández, A.M., 2017. Successful Object Encoding Induces Increased Directed Connectivity in Presymptomatic Early-Onset Alzheimer’s Disease. J Alzheimers Dis 55(3), 1195–1205.

Ossenkoppele, R., Schonhaut, D.R., Schöll, M., Lockhart, S.N., Ayakta, N., Baker, S.L., O’Neil, J.P., Janabi, M., Lazaris, A., Cantwell, A., Vogel, J., Santos, M., Miller, Z.A., Bettcher, B.M., Vossel, K.A., Kramer, J.H., Gorno-Tempini, M.L., Miller, B.L., Jagust, W.J., Rabinovici, G.D., 2016. Tau PET patterns mirror clinical and neuroanatomical variability in Alzheimer’s disease. Brain 139(Pt 5), 1551–1567.

Ossenkoppele, R., van der Flier, W.M., Zwan, M.D., Adriaanse, S.F., Boellaard, R., Windhorst, A.D., Barkhof, F., Lammertsma, A.A., Scheltens, P., van Berckel, B.N., 2013. Differential effect of APOE genotype on amyloid load and glucose metabolism in AD dementia. Neurology 80(4), 359–365.

Passamonti, L., Vázquez Rodríguez, P., Hong, Y.T., Allinson, K.S., Williamson, D., Borchert, R.J., Sami, S., Cope, T.E., Bevan-Jones, W.R., Jones, P.S., Arnold, R., Surendranathan, A., Mak, E., Su, L., Fryer, T.D., Aigbirhio, F.I., O’Brien, J.T., Rowe, J.B., 2017. 18F-AV-1451 positron emission tomography in Alzheimer’s disease and progressive supranuclear palsy. Brain 140(3), 781–791.

Poza, J., Hornero, R., Abásolo, D., Fernández, A., Mayo, A., 2008. Evaluation of spectral ratio measures from spontaneous MEG recordings in patients with Alzheimer’s disease. Comput Methods Programs Biomed 90(2), 137–147.

Rittman, T., Rubinov, M., Vértes, P.E., Patel, A.X., Ginestet, C.E., Ghosh, B.C.P., Barker, R.A., Spillantini, M.G., Bullmore, E.T., Rowe, J.B., 2016. Regional expression of the MAPT gene is associated with loss of hubs in brain networks and cognitive impairment in Parkinson disease and progressive supranuclear palsy. Neurobiol Aging 48, 153–160.

Rodriguez, E., George, N., Lachaux, J.P., Martinerie, J., Renault, B., Varela, F.J., 1999. Perception’s shadow: long-distance synchronization of human brain activity. Nature 397(6718), 430–433.

Rodriguez, R., Kallenbach, U., Singer, W., Munk, M.H., 2004. Short- and long-term effects of cholinergic modulation on gamma oscillations and response synchronization in the visual cortex. J Neurosci 24(46), 10369–10378.

Rubinov, M., Sporns, O., 2010. Complex network measures of brain connectivity: uses and interpretations. Neuroimage 52(3), 1059–1069.

Sami, S., Williams, N., Hughes, L.E., Cope, T.E., Rittman, T., Coyle-Gilchrist, I.T.S., Henson, R.N., Rowe, J.B., 2018. Neurophysiological signatures of Alzheimer’s disease and frontotemporal lobar degeneration: pathology versus phenotype. Brain 141(8), 2500–2510.

Sanz-Arigita, E.J., Schoonheim, M.M., Damoiseaux, J.S., Rombouts, S.A., Maris, E., Barkhof, F., Scheltens, P., Stam, C.J., 2010. Loss of ‘small-world’ networks in Alzheimer’s disease: graph analysis of FMRI resting-state functional connectivity. PLoS One 5(11), e13788.

Schöll, M., Lockhart, S.N., Schonhaut, D.R., O’Neil, J.P., Janabi, M., Ossenkoppele, R., Baker, S.L., Vogel, J.W., Faria, J., Schwimmer, H.D., Rabinovici, G.D., Jagust, W.J., 2016. PET Imaging of Tau Deposition in the Aging Human Brain. Neuron 89(5), 971–982.

Shankar, G.M., Bloodgood, B.L., Townsend, M., Walsh, D.M., Selkoe, D.J., Sabatini, B.L., 2007. Natural oligomers of the Alzheimer amyloid-beta protein induce reversible synapse loss by modulating an NMDA-type glutamate receptor-dependent signaling pathway. J Neurosci 27(11), 2866–2875.

Singer, W., 1999. Neuronal synchrony: a versatile code for the definition of relations? Neuron 24(1), 49-65, 111-125.

Sitnikova, T.A., Hughes, J.W., Ahlfors, S.P., Woolrich, M.W., Salat, D.H., 2018. Short timescale abnormalities in the states of spontaneous synchrony in the functional neural networks in Alzheimer’s disease. Neuroimage Clin 20, 128–152.

Smith, S.M., 2002. Fast robust automated brain extraction. Hum Brain Mapp 17(3), 143–155.

Stam, C.J., de Haan, W., Daffertshofer, A., Jones, B.F., Manshanden, I., van Cappellen van Walsum, A.M., Montez, T., Verbunt, J.P., de Munck, J.C., van Dijk, B.W., Berendse, H.W., Scheltens, P., 2009. Graph theoretical analysis of magnetoencephalographic functional connectivity in Alzheimer’s disease. Brain 132(Pt 1), 213–224.

Stam, C.J., Jones, B.F., Nolte, G., Breakspear, M., Scheltens, P., 2006. Small-world networks and functional connectivity in Alzheimer’s disease. Cereb Cortex 17(1), 92–99.

Stam, C.J., van Cappellen van Walsum, A.M., Pijnenburg, Y.A., Berendse, H.W., de Munck, J.C., Scheltens, P., van Dijk, B.W., 2002. Generalized synchronization of MEG recordings in Alzheimer’s Disease: evidence for involvement of the gamma band. J Clin Neurophysiol 19(6), 562–574.

Suarez-Revelo, J.X., Ochoa-Gomex, J.F., Duque-Grajales, J.E., Tobon-Quintero, C.A., 2016. Biomarkers identification in Alzheimer’s disease using effective connectivity analysis from electroencephalography recordings. Ingeniera e Investigacion 36, 50–57.

Supekar, K., Menon, V., Rubin, D., Musen, M., Greicius, M.D., 2008. Network analysis of intrinsic functional brain connectivity in Alzheimer’s disease. PLoS Comput Biol 4(6), e1000100.

Vecchio, F., Miraglia, F., Marra, C., Quaranta, D., Vita, M.G., Bramanti, P., Rossini, P.M., 2014. Human brain networks in cognitive decline: a graph theoretical analysis of cortical connectivity from EEG data. J Alzheimers Dis 41(1), 113–127.

Vecchio, F., Miraglia, F., Quaranta, D., Granata, G., Romanello, R., Marra, C., Bramanti, P., Rossini, P.M., 2016. Cortical connectivity and memory performance in cognitive decline: A study via graph theory from EEG data. Neuroscience 316, 143–150.

Verret, L., Mann, E.O., Hang, G.B., Barth, A.M., Cobos, I., Ho, K., Devidze, N., Masliah, E., Kreitzer, A.C., Mody, I., Mucke, L., Palop, J.J., 2012. Inhibitory interneuron deficit links altered network activity and cognitive dysfunction in Alzheimer model. Cell 149(3), 708–721.

Wang, J.H., Zuo, X.N., Gohel, S., Milham, M.P., Biswal, B.B., He, Y., 2011. Graph theoretical analysis of functional brain networks: test-retest evaluation on short- and long-term resting-state functional MRI data. PLoS One 6(7), e21976.

Zhao, X., Liu, Y., Wang, X., Liu, B., Xi, Q., Guo, Q., Jiang, H., Jiang, T., Wang, P., 2012. Disrupted small-world brain networks in moderate Alzheimer’s disease: a resting-state FMRI study. PLoS One 7(3), e33540.

